# Detecting behavioural oscillations with increased sensitivity: A modification of Brookshire’s (2022) AR-surrogate method

**DOI:** 10.1101/2024.08.22.609278

**Authors:** Anthony M. Harris, Henry A. Beale

**Affiliations:** Queensland Brain Institute, The University of Queensland, St Lucia, Australia, 4072; School of Psychology, The University of Queensland, St Lucia, Australia, 4072; School of Psychology and Neuroscience, University of Bristol, Bristol, UK, BS8 1QU; School of Health, University of the Sunshine Coast, Sippy Downs, Australia, 4556

## Abstract

A core challenge of cognitive neuroscience is to understand how cognition changes over time within individuals. For example, the tendency for behavioural responses to oscillate over time has been documented extensively in a range of cognitive domains. Recently, however, the phenomenon of behavioural oscillations has been called into question by indications that past findings might reflect aperiodic temporal structure rather than true oscillations. Brookshire (2022) proposed methods to control for aperiodic temporal structure while examining oscillations in behavioural time-courses and found no evidence of behavioural oscillations in reanalyses of four published datasets. However, Brookshire’s (2022) method has been criticised for having low sensitivity to detect effects of realistic magnitude, so it is currently unclear whether these findings suggest that behavioural oscillations are not present in these and perhaps many other datasets, or whether they are false negatives. Here, we present a modification of Brookshire’s (2022) method that uses an integrated moving average (IMA) model applied to the single-participant data, instead of the original auto-regressive (AR) approach applied to the group-level data. This method demonstrates increased sensitivity to detect effects of realistic magnitude, adequate control of false positives, and other desirable properties such as the ability to increase statistical power by adding more participants. Using this method, we reanalyse the same publicly available datasets and show significant behavioural oscillations in each of them, suggesting oscillations in behaviour are a robust phenomenon upon which to draw theoretical inferences. The participant-level IMA-surrogate method is currently the most sensitive method available for analysing behavioural oscillations while controlling for the contribution of aperiodic data fluctuations.

---

In contrast to our seemingly continuous experience of the world, a wealth of research has shown that the quality of perception and other cognitive functions fluctuates rhythmically across brief time scales (Keitel et al., 2022; Kienitz et al., 2022). For example, it has been shown that the probability of detecting a brief visual contrast transient fluctuates between better and worse detection at a rate of around 4 Hz in the second following a high-contrast ‘reset event’ (Fiebelkorn et al., 2013; Landau and Fries, 2012). Rhythmicity has been shown in a range of psychological phenomena including contrast sensitivity, phosphene perception, attentional modulation, eye-movement timing, working memory performance and many others (Dugué et al., 2016; Hogendoorn, 2016; Pomper and Ansorge, 2021; Romei et al., 2012). Such findings have been used to support influential rhythmic theories of neural processing, in which neural communication and processing are organised by rhythmic brain activity (Fries, 2023, 2015). Recently, however, criticisms have been levelled at the typical analysis methods employed in this literature. (Brookshire, 2022) argued convincingly that the typical ‘shuffling in time’ methods employed in this literature test the null hypothesis that there is no temporal structure to the data, rather than testing for the presence of rhythmicity per se. Thus, results produced by these methods may reflect aperiodic temporal structure due to alerting, inhibition of return, and so on, rather than the presence of true ‘behavioural oscillations.’ Brookshire (2022) proposed two alternative tests that directly target rhythmicity in behavioural time-courses while controlling for the influence of aperiodic trends in the data. Using these methods, Brookshire reanalysed several publicly available datasets and found no evidence for behavioural oscillations, thus casting doubt on the existence of behavioural oscillations as a phenomenon.

Interest in rhythmic properties of perception has a long and rich history in psychology and philosophy (for excellent reviews, see (Kienitz et al., 2022; Menétrey et al., 2022). Modern interest in behavioural oscillations grew out of findings that the phase of rhythmic brain activity (i.e., whether brain waves are at a peak or trough) influenced neural processing and perception (Busch et al., 2009; Dugué et al., 2011; Haegens et al., 2011; Harris et al., 2018; Mathewson et al., 2009; Williams et al., 2024). Following these findings, researchers reasoned that if neural oscillations could be aligned across different trials, such as through a phase reset, then similar phasic fluctuations should be observed in the time-course of behaviour. Early authors tested this idea by presenting a high-contrast ‘reset stimulus’ followed after a variable interval by a threshold-level target. They found that the detection rate for such targets oscillated rhythmically in the wake of a reset event (Fiebelkorn et al., 2013; Landau and Fries, 2012; Song et al., 2014), consistent with the role of neural oscillations in stimulus processing observed with EEG.

The typical approach to assessing rhythmicity in behavioural time-courses has been to compare the group-average behavioural time-course to a null distribution created by shuffling the mapping of behavioural responses to time-points to create a series of surrogate time-courses in which no temporal structure is present (Fiebelkorn et al., 2013; Landau and Fries, 2012; Tosato et al., 2022).

The amplitude spectrum produced by Fourier transforming the true behavioural data is then compared to the distribution of null spectra produced by the surrogate time-courses. The resulting p-value is the proportion of the null spectra that exceeded the amplitude of the true spectrum. This p-value is then corrected for multiple comparisons across frequencies and, if p < .05, is interpreted as evidence that oscillations are present in the pattern of behavioural data.

A concern with any application of Fourier approaches is that aperiodic (non-oscillatory) trends in the data will influence the shape of the overall amplitude spectrum, with oscillations appearing as ‘bumps’ above the general shape of this background spectrum (Donoghue et al., 2020). Recently, Brookshire (2022) pointed out that shuffling-in-time analyses remove all temporal structure from the data, both periodic and aperiodic, producing a null spectrum with roughly equal amplitude at each frequency. This ‘white noise’ null distribution is likely to differ from the amplitude spectrum produced by unshuffled data simply because there is aperiodic structure in the real data, regardless of whether or not an oscillation is present. As such, it is possible that past results produced using the shuffling-in-time method reflect aperiodic structure rather than true behavioural oscillations.

Brookshire (2022) proposed two methods to quantify the presence of behavioural oscillations while controlling for aperiodic temporal structure. Here we focus on the AR-surrogate method, as this method showed superior performance over the sampling rates and trial durations that are typical of behavioural oscillations designs. In this method, the group-averaged behavioural time-course is fit with an auto-regressive model with a single coefficient – a.k.a., an AR(1) model – that is somewhat able to capture aperiodic structure in the data but unable to model additional periodic patterns. The fitted AR(1) model is used to produce a large number of simulated behavioural time-courses that are then Fourier transformed to produce a distribution of null spectra with similar aperiodic properties to the real data.

Conceptually, the AR-surrogate approach addresses the problem of aperiodic influences that might produce false positives in the detection of behavioural oscillations. Nonetheless, this method has sparked considerable debate as it has low sensitivity to detect oscillations of realistic behavioural magnitudes (Brookshire, 2023; Fiebelkorn, 2022; Re et al., 2022; Vinck et al., 2022). For example, previous behavioural oscillation studies have shown that hit-rates in detection paradigms typically fluctuate by 15–20% during the second following a reset event (for example, hit rates might fluctuate between 65% and 85%, for a fluctuation magnitude of 20%). However, the AR-surrogate method in the form proposed and tested in (Brookshire, 2022) does not have adequate statistical power to detect simulated behavioural oscillations below a magnitude of 30%. Statistical power was even lower for behavioural-oscillation frequencies of 6 Hz and below. Thus, while theoretically appropriate, the AR-surrogate method may not be practically useful in its current form due to its propensity to give false-negative results in the presence of effects of realistic magnitude^1^.

The high false-negative rate of the AR-surrogate method when applied to simulated data creates a difficulty in the interpretation of null results when the same method is applied to real data. Brookshire (2022) re-analysed four publicly available datasets that had previously shown evidence of significant behavioural oscillations. Using those methods, non-significant results were produced for every test. Such a result might be interpreted as evidence that prior results using the shuffling-in-time method reflect aperiodic temporal structure rather than true behavioural oscillations. However, given the high false-negative rate for simulated effects of similar magnitude, the non-significant results observed with Brookshire’s methods could alternatively reflect false-negatives.

One limitation of the AR-surrogate method is that the analysis is run on the final group-averaged behavioural time-course. This means that the typical approach taken in psychology when faced with a small effect, that of testing larger groups of participants, in this case will not be very effective for increasing the power of the test. The approach we propose here is a simple modification of two aspects of Brookshire’s method that addresses this limitation. First, instead of fitting an AR(1) model, we employ an integrated moving average model of coefficient 1 (an IMA(1) model). These models are better able to produce surrogate distributions with amplitude spectra that match real data (demonstrated below). Second, instead of applying the analysis to the group-averaged time-course, we compute the IMA(1) surrogate analysis for each individual participant and then perform group-level statistical tests on the deviations of their true behavioural spectrum from their individual-level surrogate spectra. This allows us to leverage the increased statistical power for detecting small effects that results from employing larger samples. Using simulations, we show this approach increases the statistical power of the surrogate method without increasing the false positive rate at typically examined frequencies. It makes the method sensitive to behavioural oscillations of realistic magnitudes even with as few as 10 participants. It further increases the sensitivity of the analysis to behavioural oscillations in the commonly reported 3-6 Hz frequency range. Finally, we show that, when the same publicly available datasets analysed by Brookshire (2022) are re-analysed with this method, significant evidence for the existence of behavioural oscillations is obtained. Code for performing the participant-level IMA-surrogate analysis in MATLAB is available at https://github.com/AnthMHarris/BehavOsc-IMASurrogate-PLevel/.

## Methodological Details

This analysis is a simple modification of Brookshire’s (2022) approach. It uses an IMA(1) model (equivalently, an ARIMA(0,1,1) model), rather than an AR(1) to estimate the aperiodic properties of the time course, which is better at capturing the temporal properties of behavioural time courses relative to an AR(1) (**Figure 1**). The method of generating these spectra from surrogate data is described below.

**Figure 1.**
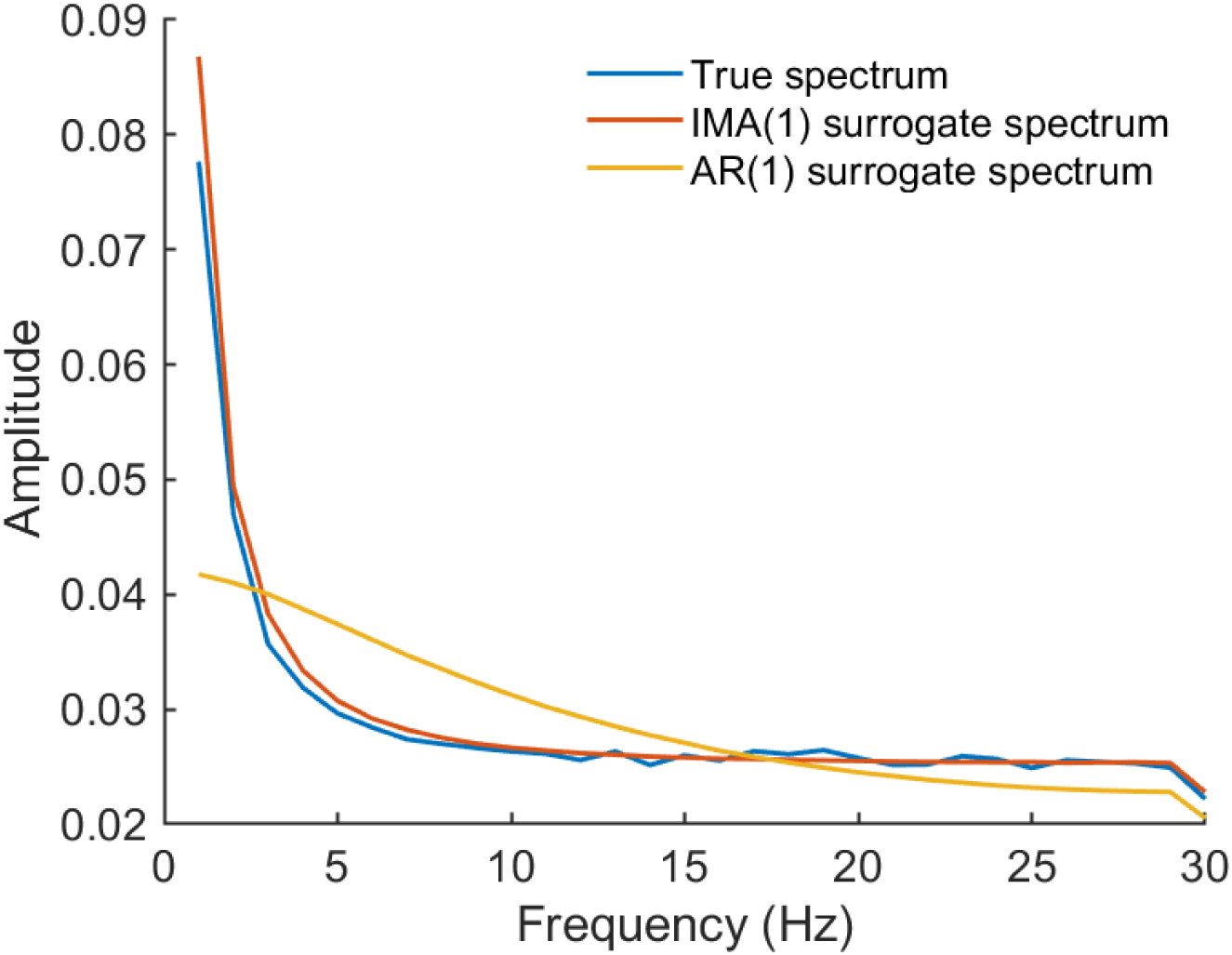
Amplitude spectra of surrogate data produced from AR(1) and IMA(1) models. The AR(1) is incapable of fully capturing the shape of the 1/f amplitude spectrum when fit with only the ∼60 samples available in a behavioural rhythms experiment. In contrast, the IMA(1) method captures the shape of the amplitude spectrum extremely well with very little data.

Additionally, our method calculates the deviation from aperiodicity for each individual, rather than for the group average (**Figure 2**). As such, this analysis addresses a subtly different question to that being asked in analyses of the group-averaged behavioural time-course. In an analysis of the group-averaged time-course, a statistical test for periodicity is asking whether significant evidence for periodicity is present in the average of participants’ behavioural reports. Such a test assumes that each individual dataset shows a similar pattern, such as an oscillation with the same frequency and phase. A test of the group-level time-course provides only indirect evidence for the presence of oscillations within the participant-level data as it is possible for patterns to emerge at the group level that are not present in any individual dataset. In comparison, applying the test at the individual level then computing group-level statistical tests asks whether, on average, evidence for rhythmicity is present directly in the participant-level data. It tests whether different individuals’ data contain rhythmicity at the same frequency, but makes no assumptions about similarity of phase angles across individuals. As these experiments are intended to assess the presence of rhythmicity in participant-level phenomena such as perception and attention, participant-level analyses are a more direct test of the theoretically relevant information.

**Figure 2.**
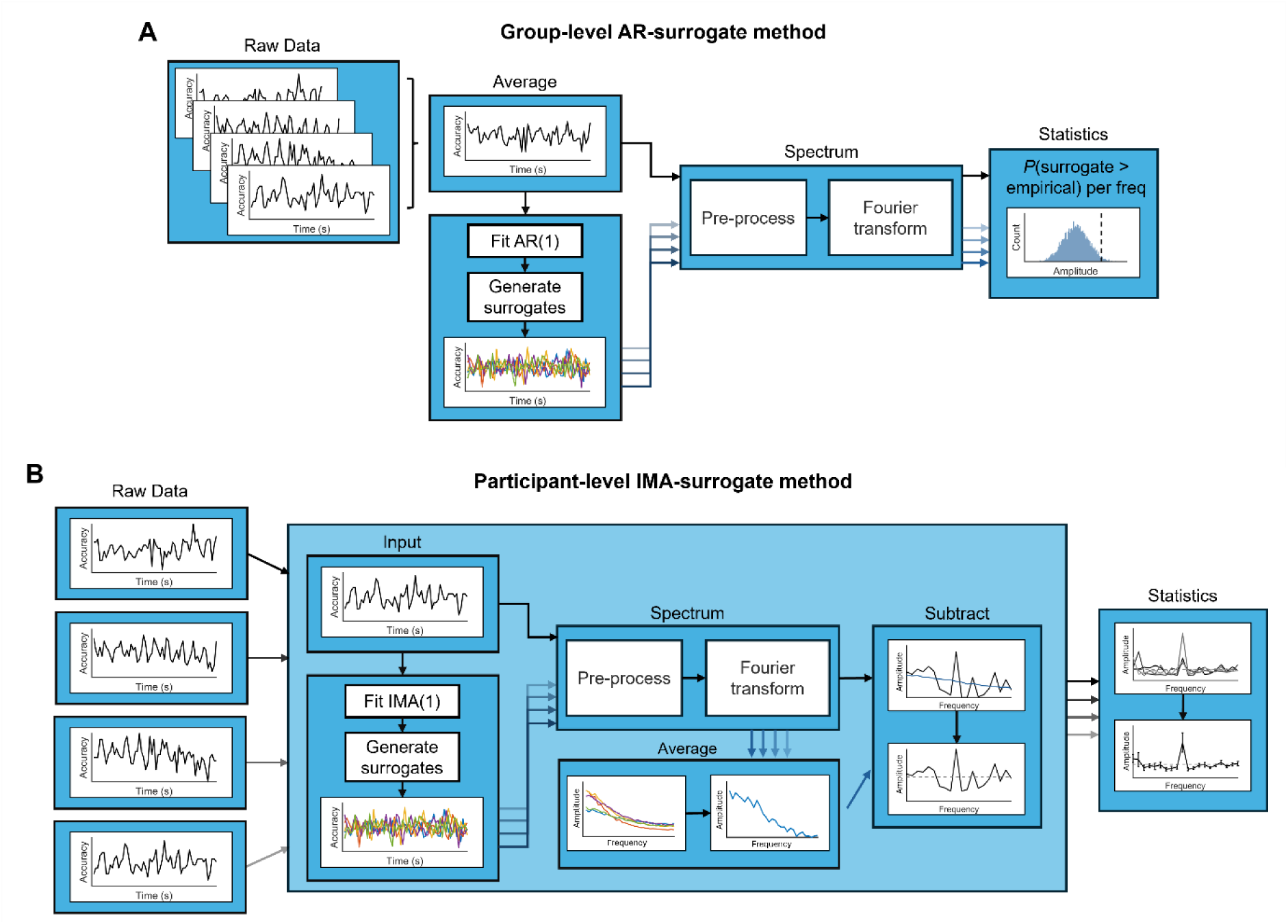
Differences between the group-level AR-surrogate and participant-level IMA-surrogate analyses. **A**) Brookshire’s (2022) group-level AR-surrogate analysis. The participant-level behavioural time-courses are averaged, and the group-average time-course is fit with an AR(1) model then used to generate a large number of surrogate time-courses. The real data and the surrogates are pre-processed in the same manner then Fourier transformed. The surrogate spectra are used to produce a null distribution of amplitude values at each frequency against which to compare the amplitude values of the real data. Typical corrections for multiple comparisons are applied (not shown here). **B**) For the participant-level IMA-surrogate analysis, each of the participant-level time-courses are fit with an IMA(1) model which is used to generate a large number of surrogate time-courses. The time-courses are pre-processed and Fourier transformed, then the average of the surrogate spectra is subtracted from the amplitude spectrum of the real data to produce a periodic spectrum. The periodic spectra for each participant are then analysed with typical statistical approaches (e.g., t-tests) at each frequency. Typical corrections for multiple comparisons are applied (not shown here). Panel A adapted from Brookshire (2022) under Creative Commons Attribution 4.0 International License: http://creativecommons.org/licenses/by/4.0/.

Once the behavioural time-course is obtained for a given participant, it is then fit with an IMA(1) model:

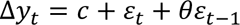

In which Δ*y_t_* is the difference in the value of the time-series at time *t* relative to time *t*-1, *c* is a constant (in this work, the constant is set to 0), *ε_t_* and *ε_t_*_−1_ are noise terms for time *t* and time *t*-1, respectively, and *θ* is the moving-average coefficient of the model that quantifies the influence of the noise at time *t-1* on the value of the moving average. This model is used to generate a large number of surrogate time-courses (e.g., 1000) with similar aperiodic properties to the original data using Monte Carlo simulation via the ‘simulate’ function in MATLAB. Any necessary preprocessing is applied identically to the true data time-course and to all of the surrogate time-courses. A method of spectral decomposition (here, the Discrete Fourier Transform) is then used to transform the true behavioural time-course into its spectral representation. The aperiodic surrogate spectrum is computed by applying the same spectral transformation to each of the surrogate time-courses and averaging the resulting surrogate spectra. This aperiodic surrogate spectrum reflects the aperiodic patterns present in the data but will not capture any oscillations that might be present. Thus, after subtracting the aperiodic surrogate spectrum from the true behavioural spectrum, we are left with a noisy representation of any periodic content in the data. We term these difference values the *periodic spectrum*, to distinguish them from the aperiodic spectrum that has been removed via subtraction.

Oscillations always appear as peaks in an amplitude spectrum relative to any aperiodic background activity. Thus, only positive values in the periodic spectrum have a meaningful interpretation. At the single-participant level, the periodic spectra are extremely noisy, making it unclear which peaks reflect genuine periodicities in the data. To determine which peaks reflect behavioural oscillations at a particular frequency, we apply standard group-level statistical approaches, to capture the variability among observations of individuals and statistically assess the between-persons reliability of the oscillation presence. As we are only interested in positive values in the periodic spectrum, we can employ one-tailed one-sample *t*-tests at each frequency, then correct for multiple comparisons with one of the many available options. Here, we employ the Bonferroni-Holm procedure (Holm, 1979). Peaks that are significant at the group level after correcting for aperiodicity in the data suggest the presence of an oscillation in the participant-level behavioural time-courses.

### Validation via Simulation

The intention of the first simulation was to test whether the proposed method can recover behavioural oscillations of a known frequency. All simulations were run using the University of Queensland’s Bunya supercomputer (The University of Queensland Research Computing Centre, 2024). We simulated participant-level data from a behavioural oscillations task with accuracy as the metric. Each participant’s data was produced (except where specified otherwise) by creating an independent idealised time-course of 1-second length, sampled at 60 Hz. This probability time-course was the sum of 1/f^2^ ‘Brownian’ noise with a 10 Hz oscillation, and was used to generate 20 random Bernoulli observations (hit = 1, miss = 0) at each time point, with the probability of a hit determined by the idealised time-course (**Figure 3A-C**). The average of observations at each timepoint produced the Response Time-course, akin to a behavioural time-course from a typical experiment (**Figure 3D**). The amplitude of the simulated oscillation corresponds to the effect size of the behavioural oscillation, which we refer to as the *behavioural depth modulation*. For example, a depth modulation of 20% indicates that the behavioural oscillation fluctuated by a difference of 20% accuracy between its peak and its trough (e.g., between 65% at the trough and 85% at the peak). We repeated this simulation at depth modulations of 0% (oscillation absent) to 50% in steps of 5%. In this simulation, the phase of the behavioural oscillation was randomly determined for each simulated participant. This procedure was performed for 500 simulated experiments at each level of depth modulation.

**Figure 3.**
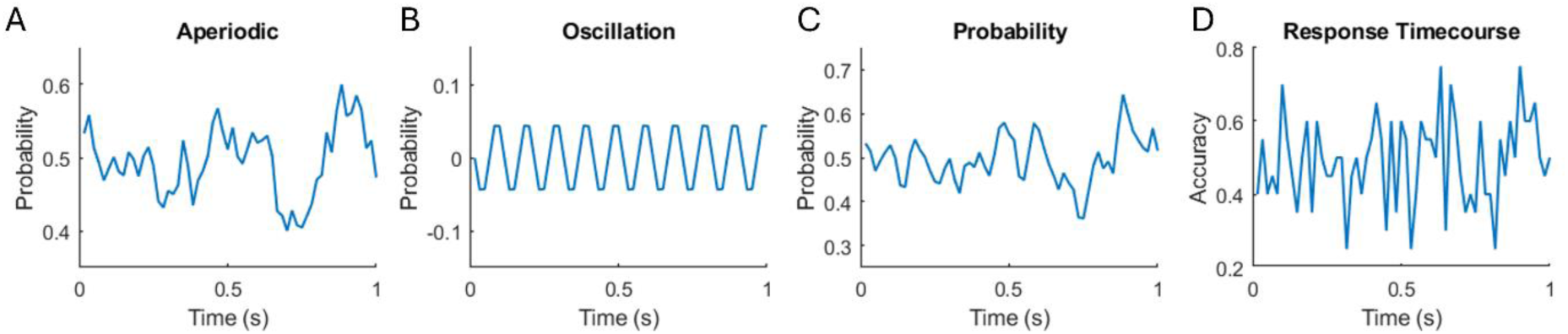
Example data simulation. The behavioural responses were simulated by creating a probability time-course (example shown in **C**) that was the sum of an aperiodic component (example shown in **A**) and an oscillatory component (example shown in **B**). After producing the probability time-course, the behavioural response time-course (**D**) was produced by comparing one random number for each simulated trial to the probability value at the relevant time-point for that trial. Values above the probability threshold were considered ‘hits’. This example reflects the process for one participant in one simulated experiment with a behavioural depth modulation of 10%.

Following the production of the simulated time-courses, we performed the analysis described above, correcting the final p-values for multiple comparisons across frequency using the Bonferroni-Holm procedure. In each simulated experiment, a significant behavioural oscillation at 10Hz is treated as a correct detection, allowing us to calculate the statistical power of such an approach. A significant oscillation detected at a depth modulation of 0% (oscillation absent) indicates a false positive and is used to calculate the false-positive rate. We also examine false positives produced at frequencies other than the simulated oscillation frequency.

The example results presented in **Figure 4** are for simulated experiments with 10 participants each. The participant-level IMA-surrogate method had high power to detect behavioural oscillations at the target frequency, 10 Hz, for all depth modulations of 15% and above. Later simulations will show how power can be increased for smaller depth modulations by increasing the sample size. The method produced very few false positives. Indeed, the false positive rate was below 5% for both the oscillation absent condition (depth modulation of 0%), and for frequencies outside 10 Hz. Thus, the method demonstrates both high sensitivity and high specificity. The following simulations are aimed at comparing the sensitivity of this method against Brookshire’s (2022) group-level AR-surrogate method and testing the robustness of these findings across different numbers of participants, trials, and across different target frequencies. It should be noted that in all cases we plot the results from the weakest combination of conditions tested (lowest participant numbers, weakest effect size, etc.) to show the limits of the method. In all cases, increasing these factors (more participants, larger effect sizes, etc.) drastically increases the statistical power of the method, and we describe these increases in the text.

**Figure 4.**
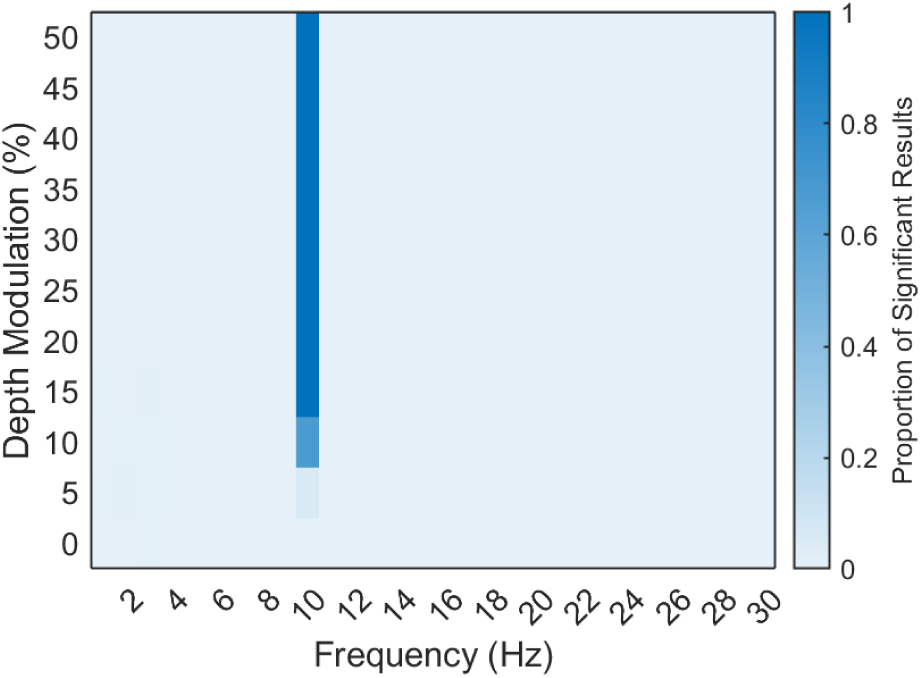
Example results for a 10 Hz behavioural oscillation in experiments with 10 participants. This simulation involved 500 experiments per depth modulation, each with 10 simulated participants. Sensitivity to detect the behavioural oscillation was above 0.95 for all depth modulations of 15% and above and false positives were low at all other frequencies. Increases in the power to detect smaller depth modulations by increasing the sample size will be shown in later simulations.

### Increasing Sensitivity by Increasing Sample Size

The purpose of this simulation was to compare the sensitivity of the participant-level IMA-surrogate method with the group-level AR-surrogate method proposed by Brookshire (2022), and to quantify the relationship between number of participants and sensitivity for the two methods. We followed the method described above to generate the simulated behavioural time-courses with a 10 Hz oscillation of varying magnitude and then analysed the same time-courses with the participant-level IMA-surrogate method and Brookshire’s (2022) group-level AR-surrogate method. We did this for each extreme of the possible behavioural oscillation space; that is, with all participants having either random or identical background aperiodic trends, and with all participants having either random or identical phases to their behavioural oscillations. When identical aperiodic trends were used, these were combined with subject-specific noise to simulate realistic inter-individual variability. As shown in **Figure 5A-D**, the participant-level IMA-surrogate method was influenced very little by whether phases were random or constant, or by the background aperiodic. When the aperiodic was random, the participant-level IMA-surrogate method achieved adequate sensitivity of 0.9 (real effects detected 90% of the time) at a depth modulation of 15% when 10 participants were simulated. With 20-150 participants, sensitivity of 0.9 was achieved at a depth modulation of 10%. For the 5% depth modulation, 80+ participants were required to achieve adequate sensitivity. When the aperiodic was identical across participants, but phase was random, the results were the same as those described above. However, sensitivity was slightly reduced when both phase and the aperiodic were identical across participants, achieving a sensitivity of 0.9 at a depth modulation of 10% with 20 simulated participants, but not achieving 0.9 sensitivity for depth modulations of 5% until sample size reached 150 participants.

**Figure 5.**
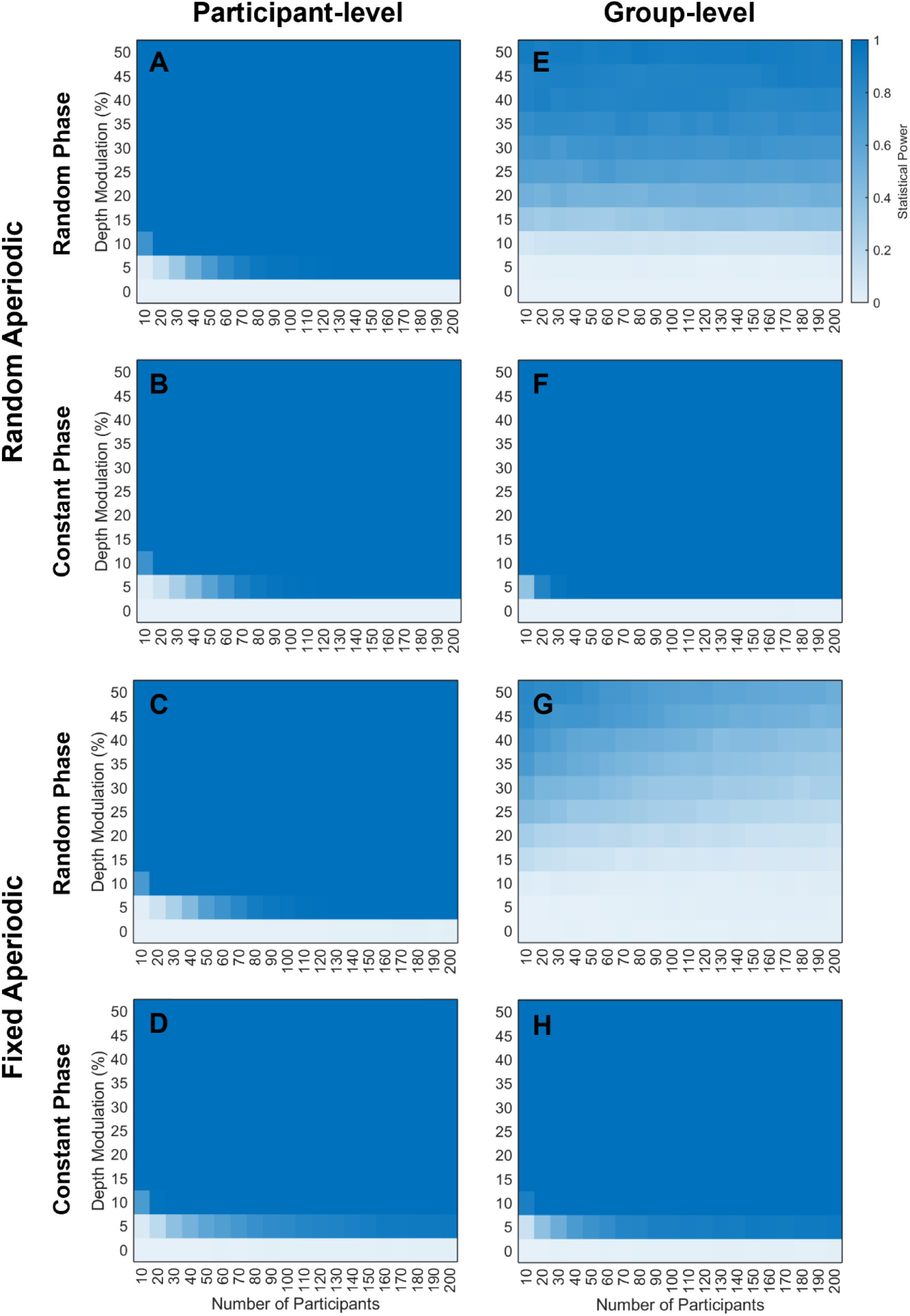
Comparison of participant-level IMA-surrogate and group-level AR-surrogate methods. The results of simulated experiments (500 per cell) analysed with the participant-level IMA-surrogate method (**A-D**) or the group-level AR-surrogate method (**E-H**). Results only shown for the target frequency (10 Hz). Across plots, we vary whether the background aperiodic is random or fixed across participants, and whether each participant’s behavioural oscillation has a different random phase or a constant phase that is shared across all participants. See headings on the left for explanations of conditions for each plot.

In contrast, the group-level AR-surrogate method (**Figure 5E-H**) was more variable across conditions. This method was weakly influenced by whether the aperiodic noise in the simulations was random or constant. However, the group-level AR-surrogate method was heavily affected by random or constant phases across participants. This is unsurprising, as the nature of the analysis is to average the time courses across participants before performing the analysis, and averaging a group of oscillations of random phase causes the oscillations to cancel out. When phases and aperiodic slopes were random across participants, this method did not reach a sensitivity of 0.9 until the participant-level depth modulation reached 50% and never reached a sensitivity of 1 within the ranges tested here. With a fixed aperiodic and random phase, this method never achieved a sensitivity of 0.9 at the depth modulations tested (note, this result is slightly different to that shown in Brookshire (2022), as the behavioural depth modulation in that paper was defined at the group level, whereas here the depth modulation is defined at the participant level). However, when phases were identical across participants, the group-level AR-surrogate method slightly outperformed the participant-level IMA-surrogate method, achieving sensitivity of 0.9 at a depth modulation of 5% with 30+ participants when the aperiodic was random, and with 100+ participants when it was constant. In each of these cases, sensitivity increased with increasing sample size, as would be expected from the noise cancellation that increasingly occurs with greater amounts of data.

These results suggest that the group-level AR-surrogate method can slightly outperform the participant-level IMA-surrogate method when phases are identical across participants but is vastly outperformed when phase is not perfectly aligned for each participant. This distinction is important, because the extent of phase alignment of behavioural oscillations across participants is currently unknown (but for evidence of considerable phase dispersion across participants, see: Liu et al., 2022; Liu and Melcher, 2023; Wang et al., 2020). Identical patterns of results across participants are extremely rare in behavioural science, particularly for such small effect sizes as the 5% depth modulation tested here. Thus, while the group-level method has the strongest performance in one extreme scenario, the participant-level IMA-surrogate method requires fewer assumptions about the data and is likely to be at least as effective as the group-level method in most realistic cases.

### Control of False Positives

To confirm that the participant-level **IMA**-surrogate method has increased sensitivity to oscillations and not simply an increased false positive rate, we analysed the proportion of experiments in which significant results were produced at frequencies with no simulated behavioural oscillation (depth modulation of 0%). For cases where aperiodic spectra were different for each participant, the proportion of significant results produced when there was no behavioural oscillation present was below .05 at all sample sizes, after controlling for multiple comparisons with the Bonferroni-Holm procedure (**Figure 6A**). This was also the case when examining the proportion of experiments producing significant results outside the simulated oscillation frequency of 10 Hz (**Figure 6B**). Indeed, false positives were below the expected level, particularly as an oscillation was introduced and the behavioural depth modulation increased (**Figure 6B**). This was due to the relative conservatism of the Bonferroni-Holm multiple-comparisons correction procedure, and because the presence of a strong oscillation in the amplitude spectrum biases the overall spectral estimate upward, reducing statistical power at other frequencies. Such a limitation could be overcome using recursive spectral estimation (Donoghue et al., 2020), but is beyond the scope of the current work.

**Figure 6.**
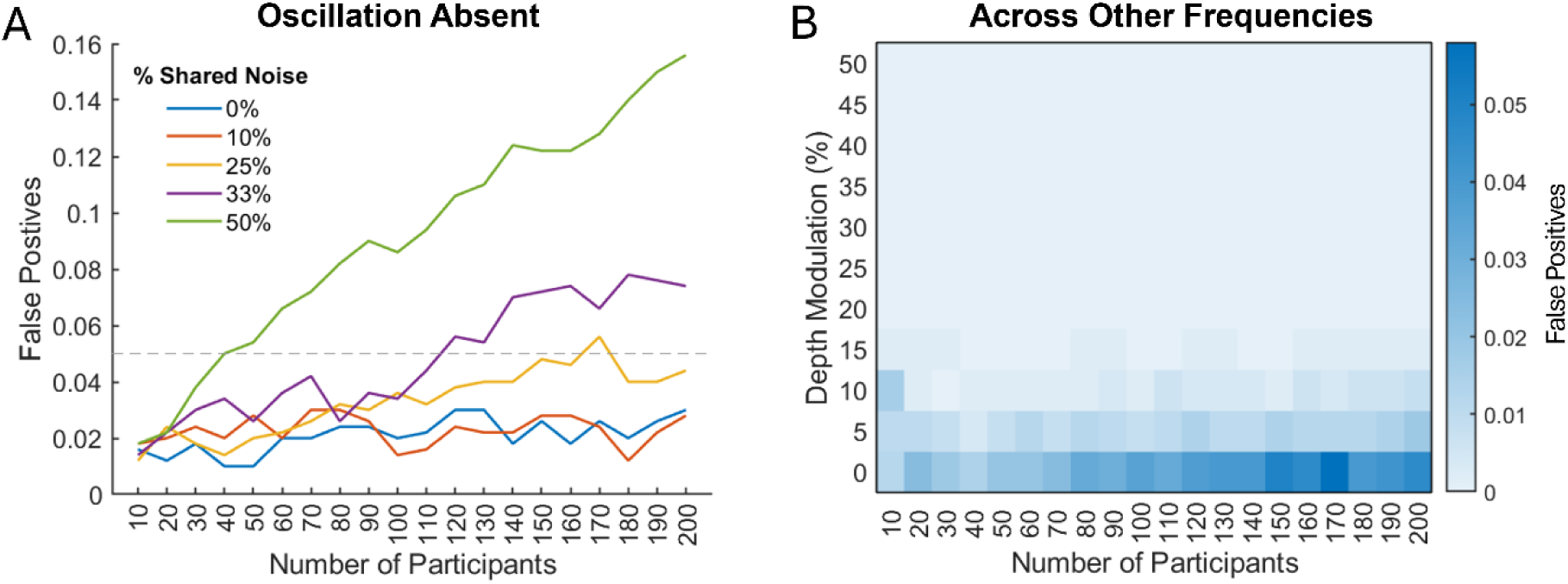
Proportion of false positives. **A**) Proportion of simulated experiments producing false positives when the behavioural oscillation was absent (Depth Modulation of 0%), across different amounts of shared group-level aperiodic structure. **B**) Proportion of simulated experiments producing significant behavioural oscillations at any frequency other than the simulated oscillation frequency across depth modulations with 25% Shared Noise. False positives decrease as depth modulation increases due to the detected oscillatory peak biasing upwards the estimated amplitude of the aperiodic background spectrum.

When the aperiodic spectra were not independent for each participant, the control of false positives depended on the extent of the shared aperiodic structure (**Figure 6A**). It is well established that consistent temporal patterns exist in behavioural data (e.g., phenomena such as Inhibition of Return; Klein, 2000), however, it is unrealistic to assume that each individual participant would show *identical* temporal structure with no individual-specific noise. Instead, we consider results from a spectrum of shared structure ranging from 0% (the above results where aperiodic spectra were unique to individuals) to 50% (identical group-level aperiodic structure makes up half of each individual’s pattern of aperiodic noise).

When shared aperiodic structure makes up 50% of the individual noise, false positives exceed .05 when 50 or more participants are used. This is because any pattern in the aperiodic activity that appears similar to an oscillation will be near-identically present in every individual due to the large amount of shared noise. With 33% shared noise, false positives emerge with around 110 participants, but for noise below this, false positives remain at or below .05 across all sample sizes. With 25% shared noise, false positives hover appropriately close to .05 at large sample sizes.

There is surprisingly little work attempting to quantify the extent of shared variance in behavioural time-courses in real human data, however, one recent study found using linear models of human reaction time data that shared noise attributable to the experimental condition explained only 2.5%-3.5% of variance in behavioural performance (Perquin et al., 2024). Adding other factors such as short- and long-term temporal dependencies across trials, blocks and sessions increased this only to 8%-17%. So, while more work is needed, it is extremely unlikely that task-related variance shared across individuals exceeds ∼25% of an individual’s behavioural noise. As such, in all the following simulations, we employ 25% shared noise across participants to estimate the statistical power of this method in the scenario of high shared noise but controlled false positives. For each subsequent analysis, the same results are shown in the Supplementary Materials for a case with no shared noise, to represent both ends of the likely extremes of real data.

### The Effect of Trial Numbers

All the simulations up to this point have been performed with 20 trials per time bin. However, many published studies have employed fewer trials than this. To examine the influence of this factor on the sensitivity of the participant-level IMA-surrogate approach, we simulated experiments with different numbers of trials per time bin. **Figure 7A** shows the statistical power to detect 10 Hz behavioural oscillations of different depth modulations with different numbers of trials per time point, specifically for a lowest sample size simulated, 10 participants. At this sample size, a statistical power of 0.9 for detecting behavioural oscillations was not achieved at a depth modulation of 10%. With 20 trials per time point, the 10% depth modulation achieved statistical power of 0.82. At depth modulations of 15%, statistical power of 0.9 was achieved with 11 trials per timepoint. For a depth modulation of 20%, only 6 trials per time point were required. As expected, the number of trials per timepoint that were required to detect an effect was reduced with increased sample sizes. With 20 participants, the number of trials required to detect behavioural oscillations of 10%, 15%, and 20% depth modulations were reduced to 12, 6, and 3, respectively.

**Figure 7.**
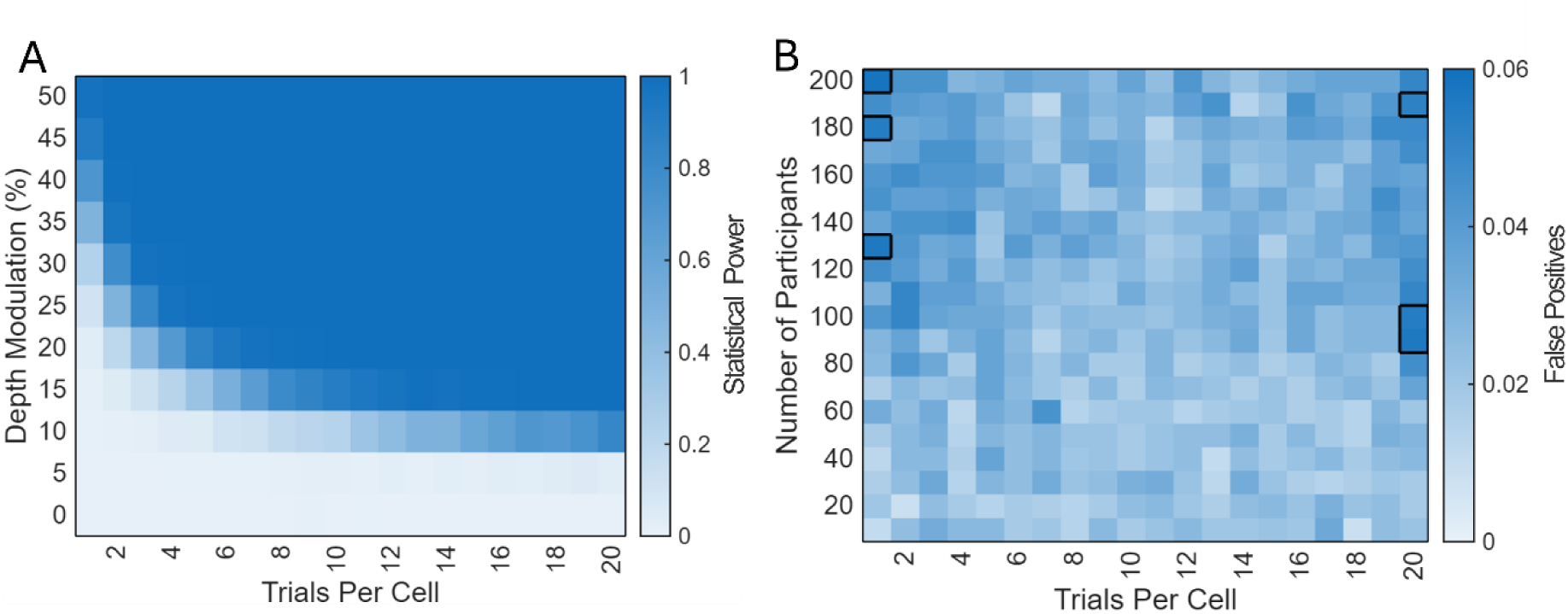
Effect of trial numbers. **A**) The sensitivity of the participant-level IMA-surrogate analysis increases with more trials-per-time-point and with increasing depth modulation. The sensitivity for a sample size of 10 participants is shown here. **B**) False positives demonstrated in the oscillation-absent condition (depth modulation 0%). Cells with solid borders indicate false positives above .05. This data was produced with 25% shared noise across participants. Refer to Figure S1 for equivalent data produced with 0% shared noise.

**Figure 7B** shows the false positive rate for different sample sizes and different numbers of trials per time-point. At all sample sizes and all trial counts, the proportion of false positives was around .05. A few cells produced average false positives up to .058, but re-running these simulations moved the cells at which these occurred, indicating they reflect stochasticity in the exact number of false alarms when simulated with a finite sample.

### Sensitivity to Oscillations of Different Frequencies

Up to this point, all the simulations have been performed with behavioural oscillations at a frequency of 10 Hz. Here, we evaluate the sensitivity of this method to detect behavioural oscillations of other relevant frequencies (with 20 trials per cell). Notably, Brookshire (2022) found with the group-level AR-surrogate method that the sensitivity to detect behavioural oscillations was reduced for oscillations at 6 Hz and below. **Figure 8A** shows the statistical power to detect behavioural oscillations of different frequencies and depth modulations, specifically for a set size of 10 participants. As this figure shows, sensitivity is affected very little by the frequency of the behavioural oscillation. **Figure 8B** shows the sensitivity to detect behavioural oscillations for different frequencies and different sample sizes at the smallest behavioural depth modulation, 5%. The statistical power to detect behavioural oscillations increased with both frequency and sample size. It should be noted, however, that this pattern was primarily present for the 5% depth modulation. For a depth modulation of 10%, statistical power was well above .9 for all frequencies of 4Hz and above at all sample sizes of 20 participants or more. Once depth modulations reach typically observed magnitudes of 15%, statistical power was at or near 1 for all frequencies between 3 Hz and 29 Hz at all sample sizes.

**Figure 8.**
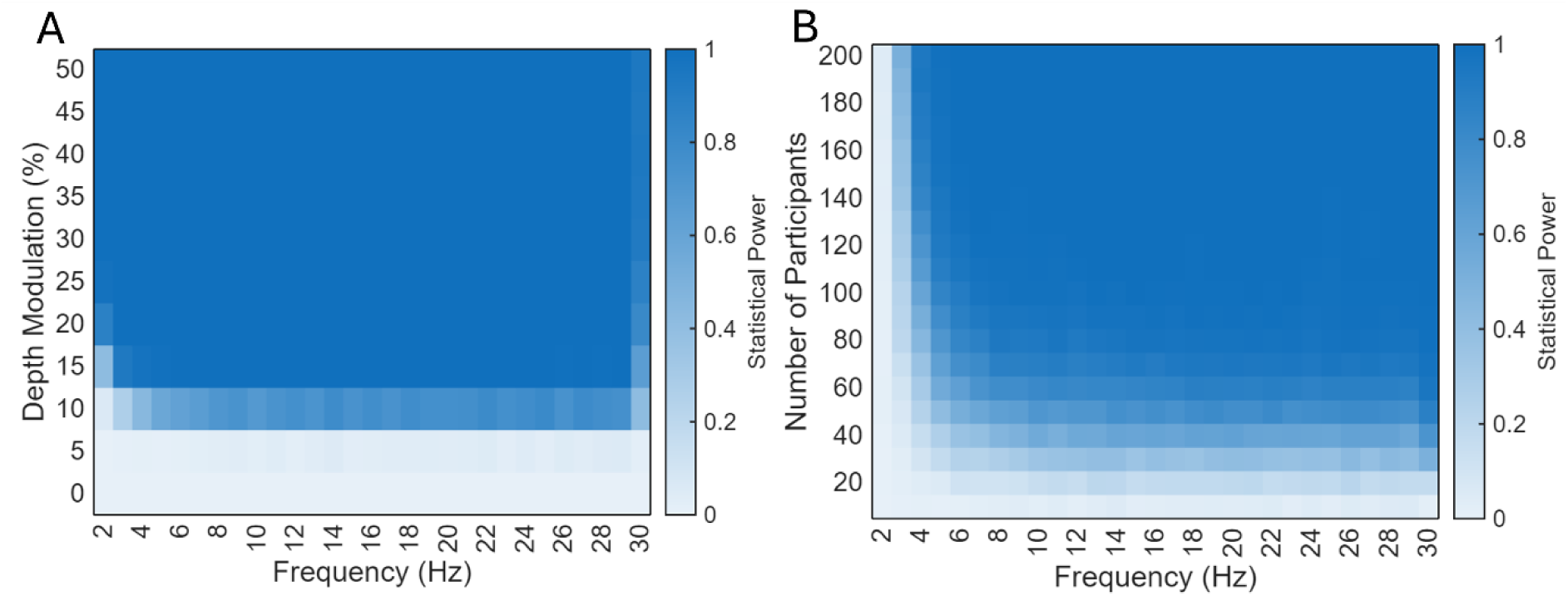
Sensitivity to oscillations of different frequencies. **A**) Target frequency by depth modulation. The example shown here is for experiments with 10 simulated participants. Statistical power is further increased at all higher participant numbers. **B**) Target frequency by number of participants. The example shown here is for the weakest depth modulation tested, 5%. For depth modulations of 10% or more, statistical power was above 0.9 at 4 Hz and higher for participant numbers of 20 or more. This data was produced with 25% shared noise across participants. Refer to Figure S2 for equivalent data produced with 0% shared noise.

### The Effect of Sampling Rate on Peak Detection at Different Frequencies

Here we examine the potential benefits of changing the sampling rate of the data for more effectively allocating limited trial numbers. We tested behavioural oscillations with sampling rates of both 30 Hz (half that tested previously) and 120 Hz (twice that tested previously). The 60 Hz simulations shown in **Figure 8** employed 20 trials per cell. To allow meaningful comparison, we simulated the 30 Hz and 120 Hz sampling rates both with the same number of trials per cell (20) and with the same number of trials in total (40 trials per cell when sampled at 30 Hz, and 10 trials per cell when sampled at 120 Hz). The results shown here are for the lowest behavioural depth modulation of 5%. It is important to note that larger effect sizes produce further statistical power improvements.

Sampling at 30 Hz with the same number of trials per cell as that shown above (20) markedly reduced statistical power across the board (**Figure 9A**). Interestingly, however, allocating the same number of total trials to a reduced sampling rate, giving 40 trials per cell at 30 Hz, increased statistical power relative to the 60 Hz sampling rate (**Figure 9C**). This was particularly apparent at low frequencies, with increases in statistical power of up to .40 at 3 Hz and up to .30 at 4 Hz. False positives were maintained below .05 in both cases (**Figure 9B&D**). Ultimately, it seems that a lower sampling rate may be useful if experimenters are specifically targeting low-frequency behavioural oscillations.

**Figure 9.**
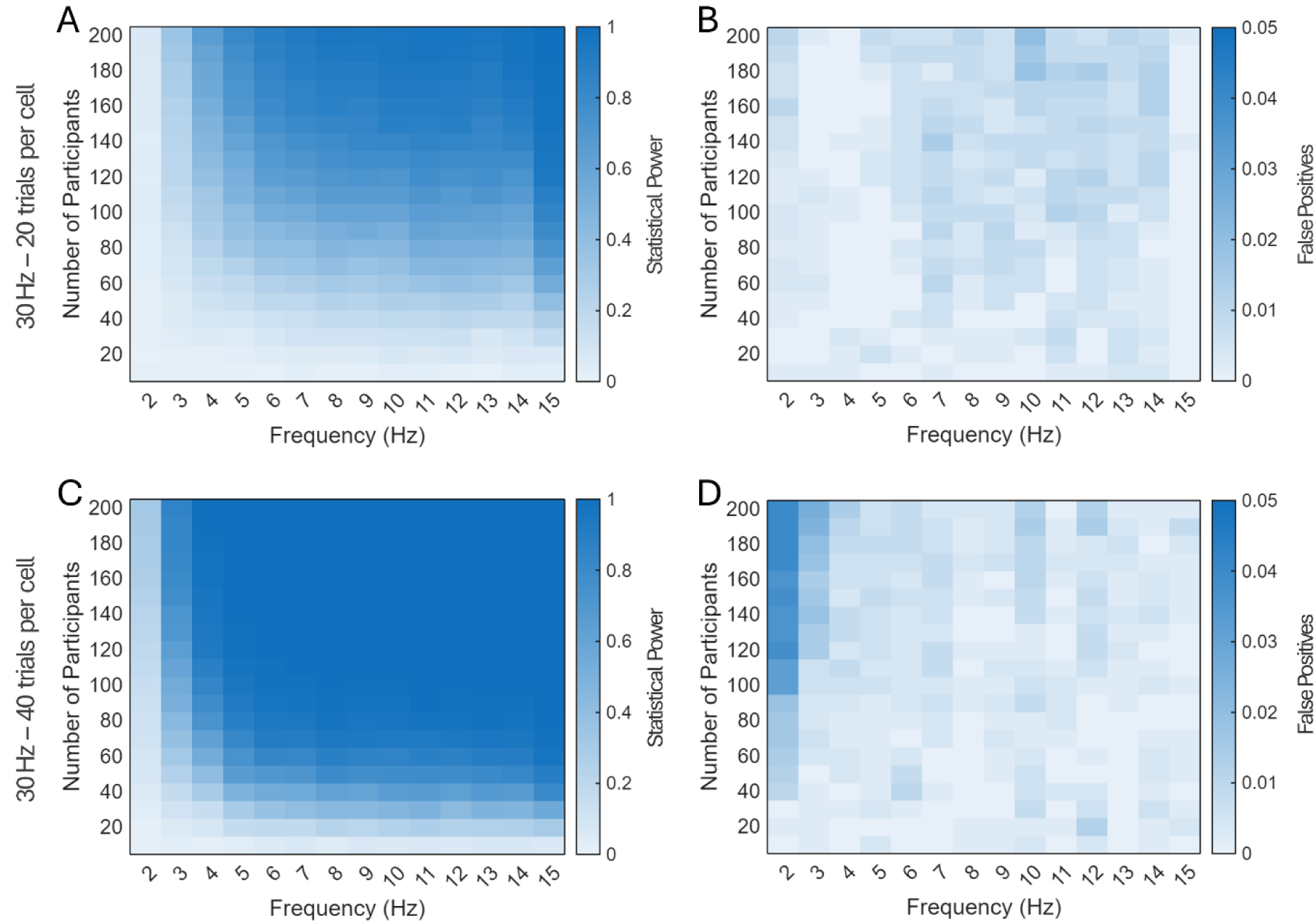
Results for a depth modulation of 5% at a sampling rate of 30 Hz. **A&B**) Results produced using 20 trials per cell, as used in Figure 8. **A**) Statistical power is generally reduced relative to the 60 Hz sampling rate. **B**) In the noise-only condition, false positives are extremely low at all frequencies and sample sizes. **C&D**) Results produced using 40 trials per cell, equivalent to the total trials per experiment in Figure 8. **C**) Statistical power is increased relative to the same number of trials divided across the 60 Hz sampling rate. This is particularly apparent at low frequencies. **D**) In the noise-only condition, false positives are below .05 at all frequencies and sample sizes. This data was produced with 25% shared noise across participants. Refer to Figure S3 for equivalent data produced with 0% shared noise.

Unsurprisingly, the 120 Hz sampling rate affected the results in the opposite direction to the 30 Hz sampling rate (**Figure 10**). Power was increased at low frequencies, relative to the 60 Hz sampling rate, when the same number of trials were used at every timepoint (doubling the number of trials per experiment), but was reduced when the trials per timepoint were halved (**Figure 10A&C**). False positives were extremely low across the board (**Figure 10B&D**). These results suggest using a higher sampling rate is only worthwhile if one is interested in resolving high-frequency behavioural oscillations.

**Figure 10.**
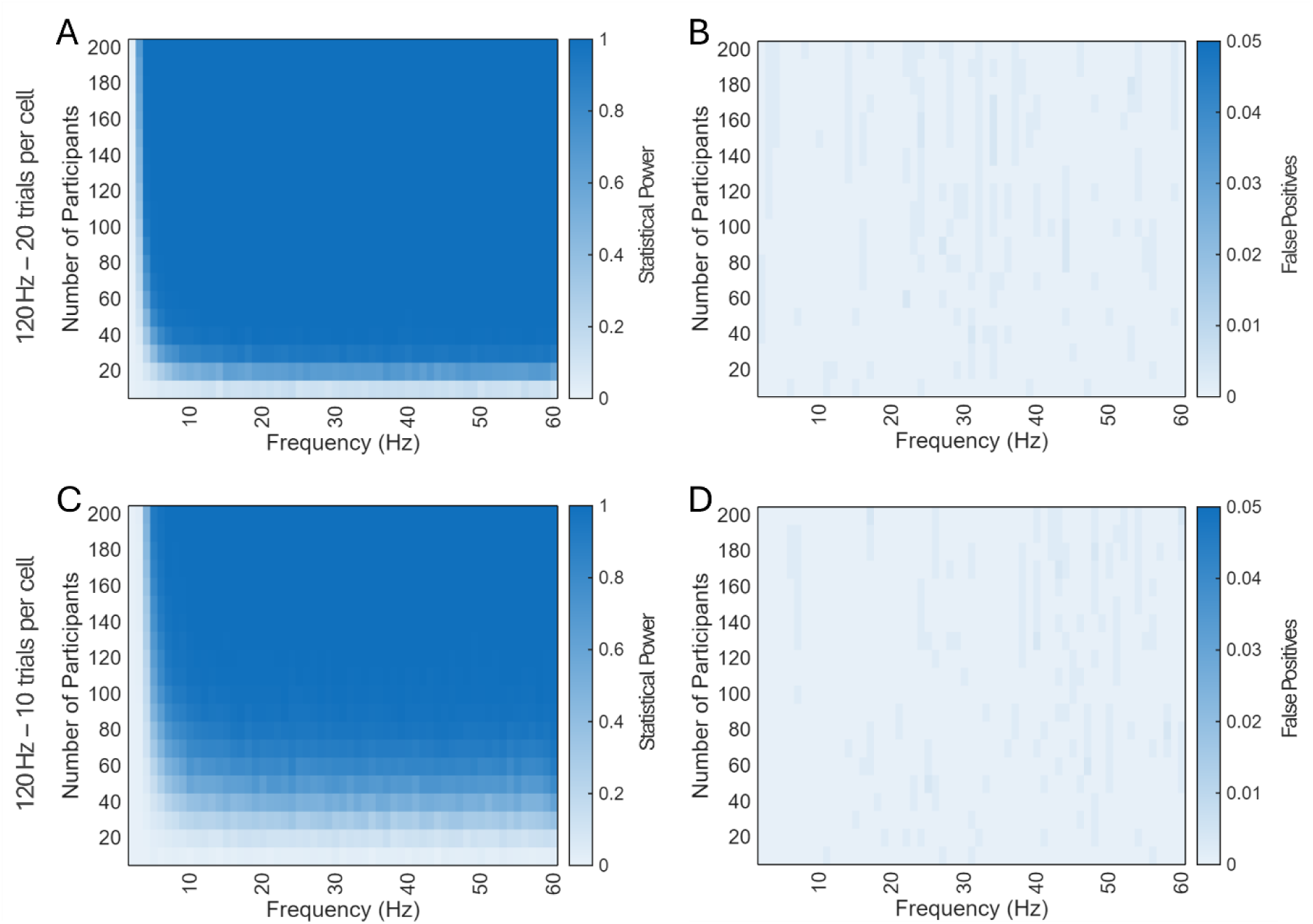
Results for a depth modulation of 5% at a sampling rate of 120 Hz. **A&B**) Results produced using 20 trials per cell, as used in Figure 8. **A**) Statistical power is improved relative to 60 Hz sampling with the same number of trials per cell. **B**) In the noise-only condition, false positives are well below .05 at all frequencies and participant numbers. **C&D**) Results produced using 10 trials per cell, equivalent to the total trials per experiment in Figure 8. **C**) Statistical power is reduced relative to the 60 Hz sampling rate. **D**) In the noise-only condition, false positives are again below .05 at all frequencies and participant numbers. This data was produced with 25% shared noise across participants. Refer to Figure S4 for equivalent data produced with 0% shared noise.

### The Effect of Detrending Behavioural Time-courses

Detrending of behavioural time-courses is common in the behavioural oscillations literature. However, Brookshire (2022) pointed out that detrending can induce spurious peaks in time-shuffled analyses, particularly when combined with tapering methods prior to Fourier analyses. Combining detrending with time course modelling can also lead to problematic distortions of the data that produce inaccurate model fits, particularly when used with short time-courses. To evaluate the influence of detrending at different stages of preprocessing on the performance of the participant-level IMA-surrogate method, we performed simulations that detrended the behavioural time-course prior to IMA(1) fitting, post-IMA(1) fitting but prior to Fourier analysis, or not at all. To further illustrate Brookshire’s (2022) point, we also performed the same procedure for the AR(1) version of our our procedure (participant-level AR(1) modelling). As shown in **Figure 12A&B**, detrending behavioural time-courses prior to IMA(1) fitting did not problematically affect the false positive rate of the method, but nor did it appreciably improve statistical power. In contrast, **Figure 12C** shows that detrending prior to AR(1) fitting had an extremely detrimental effect of the number of spurious peaks detected by the analysis. The false positive rate exceeded .05 for every sample size greater than 30 participants and grew rapidly as sample size increased. Detrending after AR(1) fitting had no detrimental effect on the number of false positives, but nor did it increase statistical power (**Figure 12D**). Based on these results, we recommend avoiding detrending when applying participant-level time-course fitting analyses as there is seemingly little benefit to be gained, and the potential for catastrophic problems in some cases.

**Figure 12.**
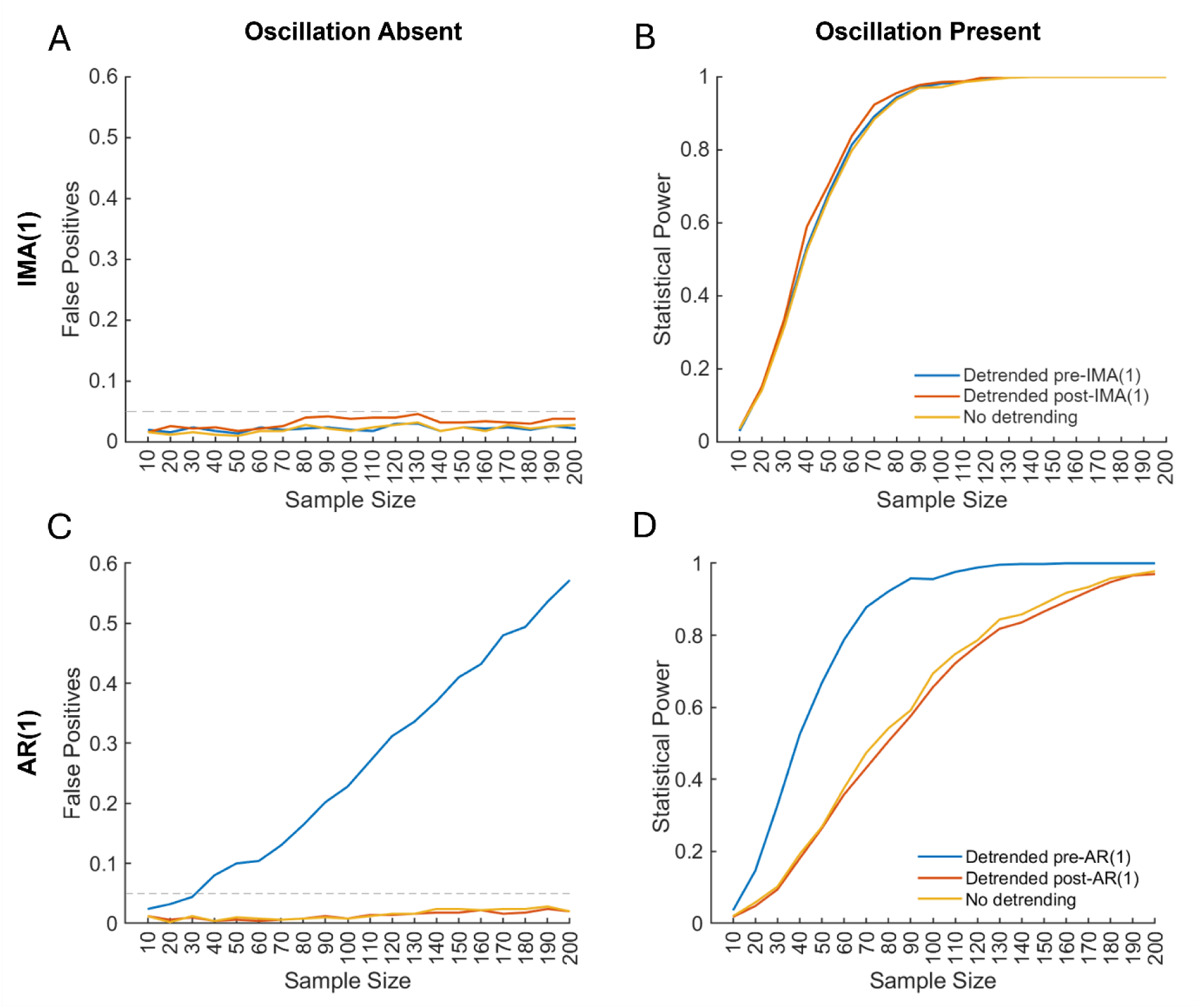
The effect of detrending on IMA(1) and AR(1) approaches to detecting behavioural oscillations. **A**) Data from the noise-only condition show that detrending has little influence on the rate of false positives when the data are modelled with an IMA(1) model. **B**) Data from the 5% depth modulation show that there is also little benefit to detrending the data when using an IMA(1) model. **C**) Data from the noise-only condition show that false positives increase dramatically with sample size when data are detrended prior to fitting AR(1) models. Detrending after AR(1) fitting has no influence on the false positive rate relative to when no detrending is used. **D**) Data from the 5% depth modulation shows that statistical power is also not meaningfully influenced by detrending after AR(1) fitting. Power appears to be higher when detrending prior to AR(1) fitting, however, this is explained by the increase in false positives shown in (A).

### Reanalyses

Brookshire (2022) reanalysed 4 publicly available datasets with the group-level AR-surrogate method and found no significant behavioural oscillations in any dataset. This raised questions about the existence of behavioural oscillations as a phenomenon, as their only supporting evidence came from shuffling-in-time methods that do not control for aperiodic patterns in the data. Here, we use the participant-level IMA-surrogate method to reanalyse 3 of the 4 datasets analysed by Brookshire (2022) and find significant behavioural oscillations in all three. The 4^th^ dataset, (Davidson et al., 2018), was not analysed here because the single-participant data was not suitable for use with this method. This is because the single-participant data did not make up smooth time-courses but was populated mostly by zero-values interspersed with sharp spikes of non-zero values that only formed a continuous time-course in the group average^2^.

It is important to reemphasize at this point that the method employed here is testing a different hypothesis to tests run on the group-average time-course that have been the norm in the past (see discussion above). As such, this is not a test of whether any specific result replicates with the participant-level IMA-surrogate analysis. Rather, the purpose of this reanalysis is to ask whether we find significant evidence supporting the existence of behavioural oscillations as a phenomenon in real behavioural time-courses. Consideration of the context of these results is outside the scope of the current work. Cursory details are given to aid comparison with the original manuscripts, but we do not provide complete information about the behavioural paradigms employed in these studies or possible interpretations of these results. The reader is referred to the original papers for further details.

Ho and colleagues (2017) reported 6 tests of behavioural oscillations in signal detection theory parameters of auditory detection, 5 of which were significant in the original paper. Reanalysis with the group-level AR-surrogate method produced no significant results (Brookshire, 2022). Re-examining this data with the participant-level IMA-surrogate method revealed 1 significant behavioural oscillation. This was an 11 Hz oscillation in the time-course of the difference in sensitivity between the two ears, *p_bh_* = .044, where *p_bh_* is the Bonferroni-Holm (1979) adjusted p-value.

Senoussi and colleagues (2019) reported 8 tests of behavioural oscillations in the time-course of visual attention, 2 of which were significant after multiple-comparison correction. Again, reanalysis with the group-level AR-surrogate method produced no significant results (Brookshire, 2022). Repeating the analyses with the participant-level IMA-surrogate method produced 1 significant result after correcting for multiple comparisons. This was an 11.5 Hz oscillation in the P1-P2 time-course for valid trials, *p_bh_* = .014.

Finally, Michel and colleagues (2021) reported 6 tests of behavioural oscillations in the time-course of the guess rate and precision parameters of mixture-modelled performance in a visual attention task. Two of these effects were significant in the original work. Reanalysis with the group-level AR-surrogate method produced no significant results (Brookshire, 2022). Repeating these tests with the current method produced 1 significant result after correction for multiple comparisons. This was a 4.8 Hz oscillation in the guess rate parameter on invalidly cued trials, *p_bh_* = .013.

## Discussion

There has been much debate recently regarding the existence of rhythmic components in the time-course of behavioural responses. While hundreds of articles exist demonstrating behavioural oscillations (for review, see Kienitz et al., 2022), they have been almost exclusively shown with analysis methods that cannot distinguish between periodic and aperiodic elements of the data (Brookshire, 2022). Brookshire (2022) proposed a method for assessing behavioural oscillations while controlling for aperiodic elements. This method was to fit an AR(1) model to the average group-level behavioural time-course and use this model to generate surrogate spectra against which to compare the amplitude spectrum of the true group-level behavioural data. Brookshire (2022) showed that several past studies produced no significant behavioural oscillations when assessed with his group-level AR-surrogate method, leading to considerable debate (Brookshire, 2023; Fiebelkorn, 2022; Re et al., 2022; Vinck et al., 2022). One interpretation of Brookshire’s (2022) results was the potential non-existence of behavioural oscillations as a phenomenon. That is, that all prior evidence of behavioural oscillations were in fact spurious results driven by aperiodic data trends. However, the method had low sensitivity to detect behavioural oscillations of magnitudes commonly observed in the literature, making the interpretation of such null results ambiguous. Here, we present a modification of the AR-surrogate approach that improves the sensitivity of the method while maintaining its adequate control of aperiodic influences. Using this participant-level IMA-surrogate method, we show in the same publicly available datasets that behavioural oscillations are indeed a robust phenomenon.

The adjustments we propose to Brookshire’s (2022) method are small: We fit a different timecourse model, an IMA(1) model, and rather than correcting for aperiodic influences in the group-average spectra, we correct them at the participant level and group-level statistical tests are applied to the resulting corrected spectra. Although the methodological changes are small, their implications are profound. Namely, that the statistical power of the test can now be increased by increasing the number of participants, allowing the method to be used to target behavioural oscillations of even small effect sizes. Additionally, the method is now robust to a number of undesirable properties that arise from examining only the group-average time-course. For example, oscillations in the group-level time-course can be produced by only one outlier participant with an unusually strong oscillation in their data, potentially producing significant results despite the pattern being absent in most participants. Alternatively, behavioural oscillations in group-level time-courses may be cancelled out if participants show behavioural oscillations with different phases. Statistical testing of the participant-level spectra addresses these problems.

A critical theoretical advantage of the participant-level IMA-surrogate method is that it takes us closer to the phenomenon that we are interested in. Tests of behavioural oscillations are intended to assess oscillations in participant-level cognitive phenomena. The assessment of these phenomena at the group level requires a number of assumptions that are not required when assessing the phenomena at the participant level. Group-level methods assume that oscillations observed in the group average are present in the data of the individuals in the group, rather than emerging as an artefact of the averaging process. They also assume that each participant shows a similar phase in their behavioural oscillation, so the oscillations will come out in the average rather than cancelling out. By using participant-level analyses we can test whether participants show evidence for behavioural oscillations without these assumptions.

Here we showed, using simulated and real data, that the participant-level IMA-surrogate method outperforms the group-level AR-surrogate method in its sensitivity to detect behavioural oscillations while maintaining adequate control of the false-positive rate. One result to highlight is that of the effect of number of participants on the sensitivity of the current method while maintaining adequate control of false positives. The group-level AR-surrogate method was highly effective at detecting oscillations that were phase locked across participants but only showed sensitivity to behavioural fluctuations of about 50% (at the participant level) when phases differed between participants. This sensitivity was not benefitted by increasing participant numbers. While some experimental data show phase-alignment across individuals (Ronconi and Melcher, 2017) many studies show substantial phase dispersion (Liu et al., 2022; Liu and Melcher, 2023; Wang et al., 2020). Without any assumptions about inter-participant phase alignment, the simulations presented here suggest that with as few as 5 trials per time-point, 20 participants are sufficient to provide >90% power to detect behavioural fluctuations of 20% and above. Performance was improved with additional participants such that >90% power to detect behavioural fluctuations of 15% was achieved with 30 participants, and for behavioural fluctuations of 10% magnitude with 60 participants completing 5 trials per time-point. These already impressive results were drastically improved by increasing the number of trials per time-point. Thus, the participant-level IMA-surrogate method is clearly sensitive to behavioural oscillations of realistic magnitudes, when assessed with realistic numbers of participants completing realistic numbers of trials.

One minor caveat is that the participant-level IMA-surrogate method only outperforms the group-level AR-surrogate method when phase is not perfectly aligned across individuals. When each individual shows a behavioural oscillation of identical phase and frequency, the noise cancellation achieved by the averaging procedure produces such a clear signal that the group-level method is able to achieve statistical power beyond what is achievable with the participant-level method. However, we also show that this benefit only matters at the smallest effect sizes, where such perfect phase alignment is most unlikely to shine through in real data, and decays quickly as we allow phase to differ between individuals. Such phase differences are likely to exist in real data for a variety of reasons, such as differences in biology that give rise to differences in conduction delays between the brain and the periphery (retina, hands, etc.), and individual differences in the peak frequency of neural oscillations (Haegens et al., 2014). Thus, while the group-level method may outperform our new method in some very specific cases, we believe the statistical gains provided by our method in the vast majority of realistic scenarios make it a valuable contribution to the field.

One troubling result of Brookshire’s (2022) work was the finding that publicly available data from past studies (Davidson et al., 2018; Ho et al., 2017; Michel et al., 2021; Senoussi et al., 2019) provided no evidence for behavioural oscillations once aperiodic contributions were controlled for. Some authors rightly pointed out that we have evidence of neural oscillations influencing behaviour that does not come from behavioural studies (Fiebelkorn, 2022), however this does not speak to the existence of oscillations in behaviour or the utility of the behavioural oscillations method of inquiry.

Behavioural oscillations provide a way to examine oscillatory contributions to processing that are difficult to examine with methods such as EEG because the oscillatory modulations occur after the effect of the stimulus-induced phase reset (Harris, 2023). Behavioural oscillations may be one of the only non-invasive methods for examining phasic fluctuations in higher-order sensory and cognitive processing. As such, the significant behavioural oscillations observed here by analysing the same publicly available datasets with the participant-level IMA-surrogate method (Ho et al., 2017; Michel et al., 2021; Senoussi et al., 2019) are exciting for two reasons. Firstly, because they reinforce the existence of behavioural oscillations as a phenomenon that reflect true oscillations in the data, rather than inadequate control of aperiodic trends. And, secondly, because they demonstrate the utility of the participant-level IMA-surrogate method in real data.

The benefit of changing from the AR-surrogate to IMA-surrogate method was demonstrated in **Figure 1**. Our testing showed that the surrogate spectra produced from an AR(1) model underestimated the exponent of the real spectra, leading to underestimated amplitudes in the tails of the spectrum and overestimated amplitudes in the centre. This had the effect of increasing false positives at low and high frequencies and reducing statistical power at intermediate frequencies. The problem persisted despite several attempts to address it with different approaches (attempts not shown here: fitting the data in loglog space, Detrended Fluctuation Analysis, and various methods for correcting the AR(1) parameter for estimation over short time series; (Donoghue et al., 2020; Kendall, 1954; Marriott and Pope, 1954; Orcutt and Winokur, 1969; Peng et al., 1994). The reality of behavioural testing with human participants means behavioural oscillations datasets typically include short time-series with low sampling rates that together produce low spectral resolution. Estimating the properties of a noisy dataset from a small sample is difficult, whether the estimation is performed using an AR(1) model or other methods (Krone et al., 2017; Thornton and Gilden, 2005; Wagenmakers et al., 2004). The IMA(1) model is more appropriate to fitting data with spectral properties typical to neural and behavioural data. It is unlikely to be a safe assumption that these features will be present in all behavioural datasets. Unfortunately, little is known about the shape of amplitude spectra derived from behavioural time-courses. No large, comprehensive review has been performed to determine their typical spectral properties, and whether these differ between measures such as accuracy, reaction time, signal-detection parameters, etc. We recommend researchers assess the noise distribution of each particular dataset and examine the adequacy of their chosen model for their particular data. It is likely that datasets with flatter spectral distributions will be better fit by an AR(1) model, and other models may be required in specific cases. The method we provide can be adjusted to incorporate any model a researcher would like to employ.

Behavioural oscillations provide critical insights into how the brain processes information across time. However, human behaviour is incredibly noisy, incorporating robust aperiodic fluctuations in addition to its periodic elements. These aperiodic fluctuations are a challenge for the assessment of behavioural oscillations. The participant-level IMA-surrogate method addresses all the problems with prior shuffling-in-time methods while providing good sensitivity to detecting even small oscillatory effects. We hope it will be of use in the study of rhythmic processes in cognition.

## Code Availability

Code for performing the participant-level IMA-surrogate analysis is available at https://github.com/AnthMHarris/BehavOsc-IMASurrogate-PLevel/

## Footnotes

1 Of course, careful experimental design is also required to ameliorate some of the effects of measurement noise on the detectability of smaller effects (Brookshire, 2023).

2 Thanks to Matt Davidson for providing the single-participant data for their study so this could be assessed.

## References

Brookshire G. 2023. Distinguishing between periodicity and consistency over trials: Reply to Re, Tosato, Fries, and Landau. DOI: 10.31234/osf.io/km7e9

Brookshire G. 2022. Putative rhythms in attentional switching can be explained by aperiodic temporal structure. Nature Human Behaviour 1–12. DOI: 10.1038/s41562-022-01364-0, PMID: 35680992

Busch NA, Dubois J, VanRullen R. 2009. The Phase of Ongoing EEG Oscillations Predicts Visual Perception. The Journal of Neuroscience 29:7869–7876. DOI: 10.1523/jneurosci.0113-09.2009, PMID: 19535598

The University of Queensland Research Computing Centre. 2024. Bunya supercomputer. DOI: 10.48610/wf6c-qy55

Davidson MJ, Alais D, Boxtel JJ van, Tsuchiya N. 2018. Attention periodically samples competing stimuli during binocular rivalry. eLife 7:e40868. DOI: 10.7554/elife.40868, PMID: 30507378

Donoghue T, Haller M, Peterson EJ, Varma P, Sebastian P, Gao R, Noto T, Lara AH, Wallis JD, Knight RT, Shestyuk A, Voytek B. 2020. Parameterizing neural power spectra into periodic and aperiodic components. Nature Neuroscience 23:1655–1665. DOI: 10.1038/s41593-020-00744-x

Dugué L, Marque P, VanRullen R. 2011. The Phase of Ongoing Oscillations Mediates the Causal Relation between Brain Excitation and Visual Perception. The Journal of Neuroscience 31:11889– 11893. DOI: 10.1523/jneurosci.1161-11.2011, PMID: 21849549

Dugué L, Roberts M, Carrasco M. 2016. Attention Reorients Periodically. Current Biology 26:1595–1601. DOI: 10.1016/j.cub.2016.04.046, PMID: 27265395

Fiebelkorn IC. 2022. There Is More Evidence of Rhythmic Attention than Can Be Found in Behavioral Studies: Perspective on Brookshire,. Journal of Cognitive Neuroscience 35:128–134. DOI: 10.1162/jocn_a_01936, PMID: 36306250

Fiebelkorn IC, Saalmann YB, Kastner S. 2013. Rhythmic Sampling within and between Objects despite Sustained Attention at a Cued Location. Current Biology 23:2553–2558. DOI: 10.1016/j.cub.2013.10.063, PMID: 24316204

Fries P. 2023. Rhythmic attentional scanning. Neuron 111:954–970. DOI: 10.1016/j.neuron.2023.02.015, PMID: 37023718

Fries P. 2015. Rhythms for Cognition: Communication through Coherence. Neuron 88:220–235. DOI: 10.1016/j.neuron.2015.09.034, PMID: 26447583

Haegens S, Cousijn H, Wallis G, Harrison PJ, Nobre AC. 2014. Inter- and intra-individual variability in alpha peak frequency. NeuroImage 92:46–55. DOI: 10.1016/j.neuroimage.2014.01.049, PMID: 24508648

Haegens S, Nácher V, Luna R, Romo R, Jensen O. 2011. α-Oscillations in the monkey sensorimotor network influence discrimination performance by rhythmical inhibition of neuronal spiking. Proceedings of the National Academy of Sciences 108:19377–19382. DOI: 10.1073/pnas.1117190108, PMID: 22084106

Harris AM. 2023. Phase resets undermine measures of phase-dependent perception. Trends in Cognitive Sciences 27:224–226. DOI: 10.1016/j.tics.2022.12.008, PMID: 36609017

Harris AM, Dux PE, Mattingley JB. 2018. Detecting Unattended Stimuli Depends on the Phase of Prestimulus Neural Oscillations. Journal of Neuroscience 38:3092–3101. DOI: 10.1523/jneurosci.3006-17.2018, PMID: 29459372

Ho H, Leung J, Burr DC, Alais D, Morrone M. 2017. Auditory Sensitivity and Decision Criteria Oscillate at Different Frequencies Separately for the Two Ears. Current Biology. DOI: 10.1016/j.cub.2017.10.017

Hogendoorn H. 2016. Voluntary saccadic eye movements ride the attentional rhythm. Journal of cognitive neuroscience 28:1625–1635.

Holm S. 1979. A Simple Sequentially Rejective Multiple Test Procedure. Scandanavian Journal of Statistics 6:65–70.

Keitel C, Ruzzoli M, Dugué L, Busch NA, Benwell CSY. 2022. Rhythms in Cognition: The evidence revisited. European Journal of Neuroscience. DOI: 10.1111/ejn.15740, PMID: 35696729

Kendall MG. 1954. Note on bias in the estimation of autocorrelation. Biometrika 41:403–404. DOI: 10.1093/biomet/41.3-4.403

Kienitz R, Schmid MC, Dugué L. 2022. Rhythmic sampling revisited: Experimental paradigms and neural mechanisms. European Journal of Neuroscience 55:3010–3024. DOI: 10.1111/ejn.15489, PMID: 34643973

Krone T, Albers CJ, Timmerman ME. 2017. A comparative simulation study of AR(1) estimators in short time series. Quality & Quantity 51:1–21. DOI: 10.1007/s11135-015-0290-1, PMID: 28133396

Landau A, Fries P. 2012. Attention Samples Stimuli Rhythmically. Current Biology 22:1000–1004. DOI: 10.1016/j.cub.2012.03.054, PMID: 22633805

Liu X, Balestrieri E, Melcher D. 2022. Evidence for a theta-band behavioural oscillation in rapid face detection. European Journal of Neuroscience 56:5033–5046. DOI: 10.1111/ejn.15790, PMID: 35943892

Liu X, Melcher D. 2023. The effect of familiarity on behavioral oscillations in face perception. Scientific Reports 13:10145. DOI: 10.1038/s41598-023-34812-6, PMID: 37349366

Marriott FHC, Pope JA. 1954. Bias in the Estimation of Autocorrelations. Biometrika 41:390–402.

Mathewson KE, Gratton G, Fabiani M, Beck DM, Ro T. 2009. To See or Not to See: Prestimulus α Phase Predicts Visual Awareness. The Journal of Neuroscience 29:2725–2732. DOI: 10.1523/jneurosci.3963-08.2009, PMID: 19261866

Menétrey MQ, Vogelsang L, Herzog MH. 2022. A guideline for linking brain wave findings to the various aspects of discrete perception. European Journal of Neuroscience 55:3528–3537. DOI: 10.1111/ejn.15349, PMID: 34125452

Michel R, Dugué L, Busch NA. 2021. Distinct contributions of alpha and theta rhythms to perceptual and attentional sampling. European Journal of Neuroscience. DOI: 10.1111/ejn.15154, PMID: 33609313

Orcutt GH, Winokur HS. 1969. First Order Autoregression: Inference, Estimation, and Prediction. Econometrica 37:1. DOI: 10.2307/1909199

Peng C-K, Buldyrev SV, Havlin S, Simons M, Stanley HE, Goldberger AL. 1994. Mosaic organization of DNA nucleotides. Physical Review E 49:1685–1689. DOI: 10.1103/physreve.49.1685, PMID: 9961383

Perquin MN, Heed T, Kayser C. 2024. Variance (Un)Explained: Experimental Conditions and Temporal Dependencies Explain Similarly Small Proportions of Reaction Time Variability in Linear Models of Perceptual and Cognitive Tasks. Journal of Experimental Psychology: General 153:3107–3129. DOI: 10.1037/xge0001630, PMID: 39311860

Pomper U, Ansorge U. 2021. Theta-Rhythmic Oscillation of Working Memory Performance. Psychological science 32:1801–1810. DOI: 10.1177/09567976211013045, PMID: 34592108

Re D, Tosato T, Fries P, Landau AN. 2022. Perplexity about periodicity repeats perpetually: A response to Brookshire. bioRxiv 2022.09.26.509017. DOI: 10.1101/2022.09.26.509017

Romei V, Gross J, Thut G. 2012. Sounds Reset Rhythms of Visual Cortex and Corresponding Human Visual Perception. Current Biology 22:807–813. DOI: 10.1016/j.cub.2012.03.025, PMID: 22503499

Ronconi L, Melcher D. 2017. The Role of Oscillatory Phase in Determining the Temporal Organization of Perception: Evidence from Sensory Entrainment. The Journal of Neuroscience 37:10636– 10644. DOI: 10.1523/jneurosci.1704-17.2017, PMID: 28972130

Senoussi M, Moreland JC, Busch NA, Dugué L. 2019. Attention explores space periodically at the theta frequency. Journal of Vision 19:22. DOI: 10.1167/19.5.22, PMID: 31121012

Song K, Meng M, Chen L, Zhou K, Luo H. 2014. Behavioral Oscillations in Attention: Rhythmic α Pulses Mediated through θ Band. The Journal of Neuroscience 34:4837–4844. DOI: 10.1523/jneurosci.4856-13.2014, PMID: 24695703

Thornton TL, Gilden DL. 2005. Provenance of correlations in psychological data. Psychonomic Bulletin & Review 12:409–441. DOI: 10.3758/bf03193785, PMID: 16235626

Tosato T, Rohenkohl G, Dowdall JR, Fries P. 2022. Quantifying rhythmicity in perceptual reports. NeuroImage 262:119561. DOI: 10.1016/j.neuroimage.2022.119561, PMID: 35973565

Vinck M, Uran C, Schneider M. 2022. Aperiodic processes explaining rhythms in behavior: A matter of false detection or definition? DOI: 10.31234/osf.io/wzvfh

Wagenmakers E-J, Farrell S, Ratcliff R. 2004. Estimation and interpretation of 1/fα noise in human cognition. Psychonomic Bulletin & Review 11:579–615. DOI: 10.3758/bf03196615, PMID: 15581115

Wang M, Huang Y, Luo H, Zhang H. 2020. Sustained Visual Priming Effects Can Emerge from Attentional Oscillation and Temporal Expectation. The Journal of Neuroscience 40:3657–3674. DOI: 10.1523/jneurosci.2539-19.2020, PMID: 32253359

Williams JG, Harrison WJ, Beale HA, Mattingley JB, Harris AM. 2024. Effects of neural oscillation power and phase on discrimination performance in a visual tilt illusion. Current Biology. DOI: 10.1016/j.cub.2024.03.014

