## Supplementary Materials for "Detecting behavioural oscillations with increased sensitivity: A modification of Brookshire’s (2022) AR-surrogate method"

A.M. Harris & H.A. Beale

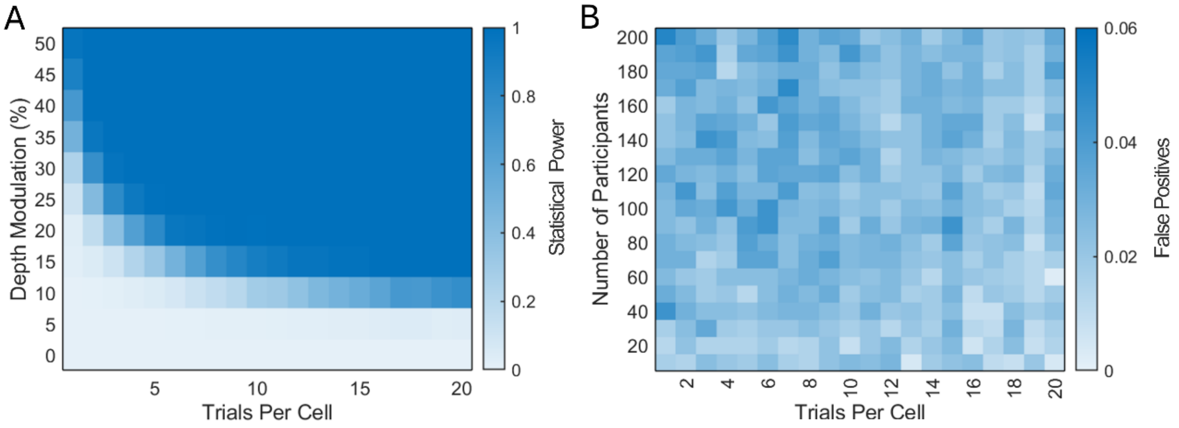


**Figure S1. Effect of trial numbers with no shared noise. A**) The sensitivity of the participant-level IMA-surrogate analysis increases with more trials-per-time-point and with increasing depth modulation. The sensitivity for a sample size of 10 participants is shown here. **B**) False positives demonstrated in the oscillation-absent condition (depth modulation 0%). Cells with solid borders indicate false positives above .05 (there were none here). Data are the equivalent of Figure 7 in the main paper with no shared noise across participants.

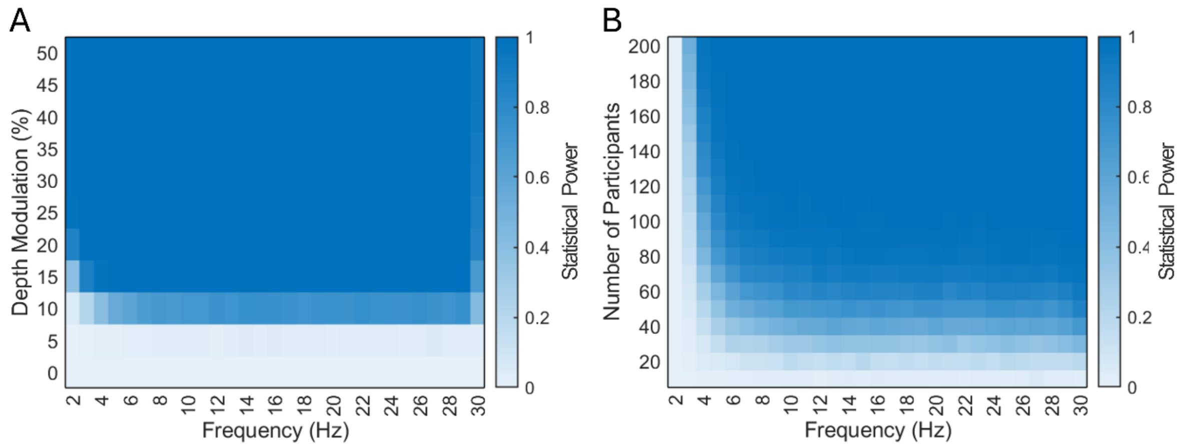


**Figure S2. Sensitivity to oscillations of different frequencies with no shared noise. A**) Target frequency by depth modulation. The example shown here is for experiments with 10 simulated participants. Statistical power is further increased at all higher participant numbers. **B**) Target frequency by number of participants. The example shown here is for the weakest depth modulation tested, 5%. For depth modulations of 10% or more, statistical power was above 0.9 at 4 Hz and higher for participant numbers of 20 or more. Data are the equivalent of Figure 8 in the main paper with no shared noise across participants.

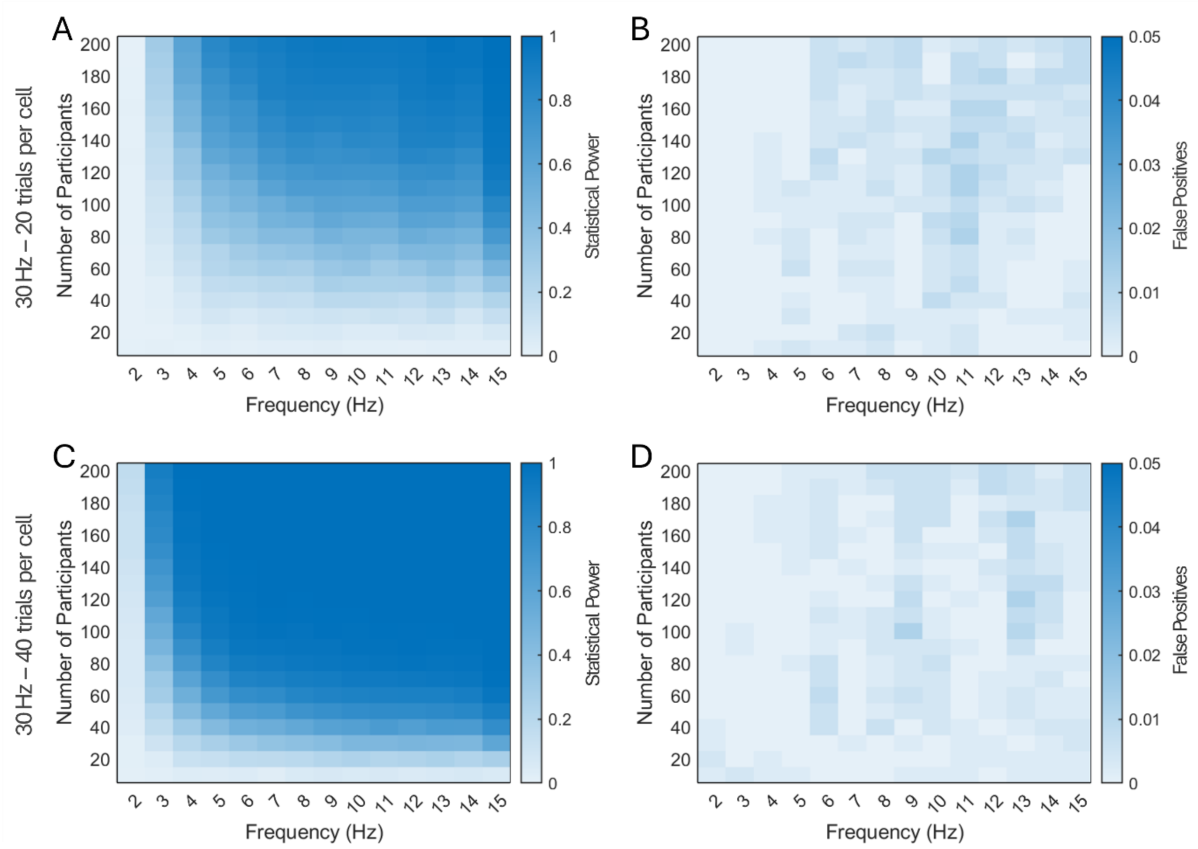


**Figure S3. Results for a depth modulation of 5% at a sampling rate of 30 Hz with no shared noise. A&B**) Results produced using 20 trials per cell, as used in Figure 8. **A**) Statistical power is generally reduced relative to the 60 Hz sampling rate. **B**) In the noise-only condition, false positives are extremely low at all frequencies and sample sizes. **C&D**) Results produced using 40 trials per cell, equivalent to the total trials per experiment in Figure 8. **C**) Statistical power is increased relative to the same number of trials divided across the 60 Hz sampling rate. This is particularly apparent at low frequencies. **D**) In the noise-only condition, false positives are below .05 at all frequencies and sample sizes. Data are the equivalent of Figure 9 in the main paper with no shared noise across participants.

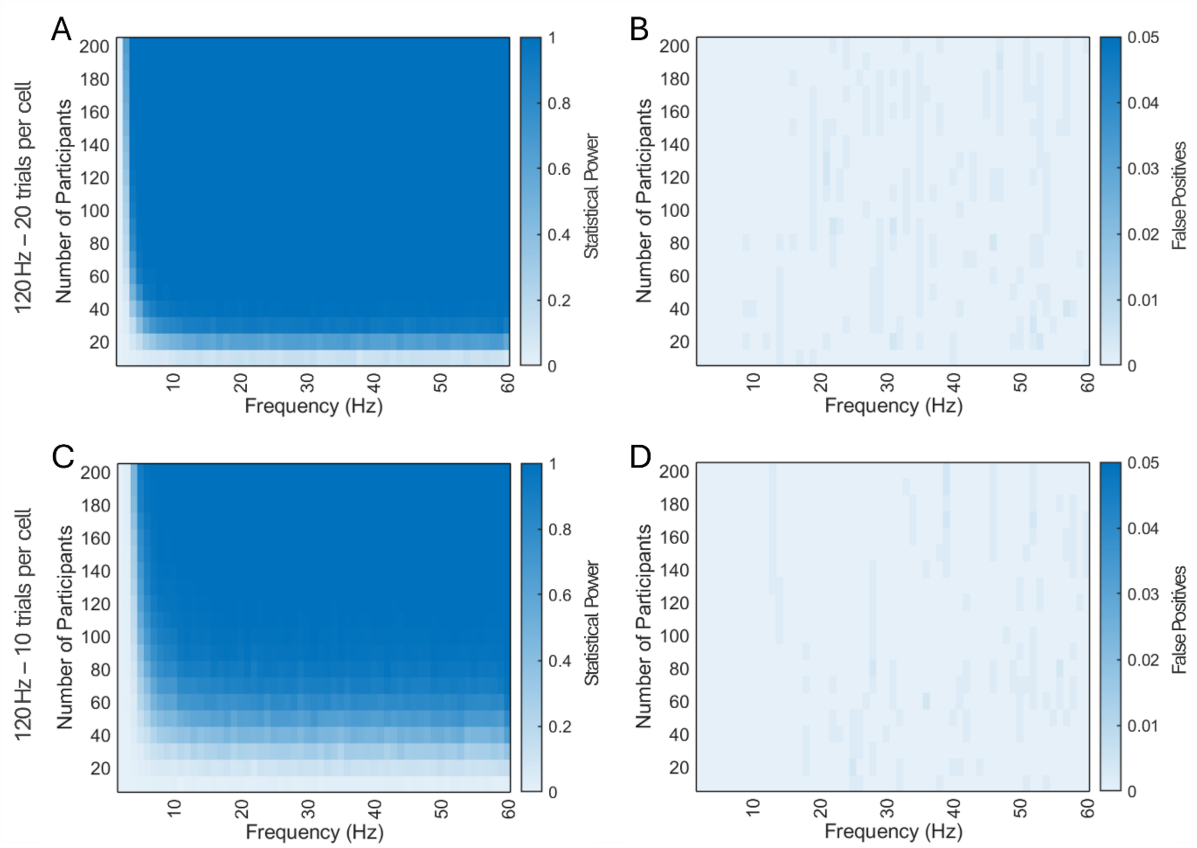


**Figure S4. Results for a depth modulation of 5% at a sampling rate of 120 Hz with no shared noise. A&B**) Results produced using 20 trials per cell, as used in Figure 8. **A**) Statistical power is improved relative to 60 Hz sampling with the same number of trials per cell. **B**) In the noise-only condition, false positives are well below .05 at all frequencies and participant numbers. **C&D**) Results produced using 10 trials per cell, equivalent to the total trials per experiment in Figure 8. **C**) Statistical power is reduced relative to the 60 Hz sampling rate. **D**) In the noise-only condition, false positives are again below .05 at all frequencies and participant numbers. Data are the equivalent of Figure 10 in the main paper with no shared noise across participants.
